# Heterogeneity of burst firing in mouse thalamic reticular nucleus neurons

**DOI:** 10.1101/2022.09.25.509400

**Authors:** Laura M. Harding-Jackson, Joseph A. Beatty, Charles L. Cox

## Abstract

The thalamic reticular nucleus (TRN) sits at the interface of the thalamus and neocortex and provides the majority of inhibition to thalamic relay nuclei. Functionally, the activity of TRN neurons can impact sensory processing and may influence arousal states. TRN neurons discharge action potentials in two distinct output modes: tonic or burst firing. Burst output, a transient high frequency discharge of action potentials, is dependent on the activation of transient low-threshold voltage-dependent T-type calcium current (I_T_). In our current study, we identify a broad range of burst firing frequencies in TRN neurons, which depend on the activation of I_T_. The amplitude of the low-threshold calcium spike (LTS) underlying the burst positively correlated with burst frequency and number of action potentials per burst. Activation of small conductance calcium-activated potassium (SK) channels on TRN neurons can impact burst discharge. Attenuation of SK channels increased TRN neuron burst frequency through an increase in LTS duration, but not magnitude. The broad range of burst firing frequencies could provide distinct downstream inhibition within thalamic nuclei.

## INTRODUCTION

The thalamic reticular nucleus (TRN) serves as a major source of inhibition to most thalamic nuclei ^1–3^. The TRN forms a shell around the dorsal thalamus, positioning it at the interface between the neocortex and thalamus. In the sensory systems, TRN neurons receive excitatory synaptic input from ascending thalamocortical efferents and descending corticothalamic efferents ^4,5^. TRN neurons integrate this afferent activity, provide inhibitory output to thalamic nuclei, thereby modulating thalamocortical circuit activity. The modulatory impact of TRN is not limited to only sensory systems, but can impact motor and limbic systems as well ^6,7^.

TRN neurons, like thalamocortical neurons, produce two distinct modes of action potential output: burst and tonic. Burst mode is typically characterized by a transient, high frequency discharge of action potentials (>250 Hz) requiring the activation of low-threshold transient calcium (T) channels ^8–13^. At relatively hyperpolarized potentials, activation of T-channels gives rise to a large, transient membrane depolarization, referred to as a low-threshold spike (LTS). The LTS is crowned by a high frequency discharge of multiple action potentials. At relatively depolarized membrane potentials, T-channels are inactivated and suprathreshold membrane depolarizations trigger an action potential discharge that is linearly related to the magnitude of the depolarization referred to as tonic discharge ^14^. Rhythmic burst discharge has been associated with specific behavioral states such as slow-wave sleep and certain seizure disorders ^15,16^. Thalamic burst discharge during awake states has been observed with changes in attentional state or locomotion ^8,17–19^. Burst and tonic firing can have significantly different impacts on downstream thalamocortical neurons. Burst discharge of TRN neurons produces long-lasting inhibitory responses dependent on the activation of both GABA_A_ and GABA_B_ receptors. In contrast, tonic discharge predominantly produces short duration inhibition dependent on GABA_A_ receptor activation ^3,20,21^.

Burst magnitude, duration, and rhythmicity can be modulated by a variety of currents intrinsic to thalamic reticular neurons. Small conductance calcium-activated potassium (SK) channels have been shown to decrease the intensity of the burst discharge ^22^. Rhythmic oscillations in TRN neurons are maintained by the interplay of selective sarcoplasmic reticulum calcium ATPase (SERCA) and SK channels, which rapidly uptake calcium to allow T-channels to quickly recover ^23–25^. It is clear from these studies that SK-channel activation has a prominent role in shaping the burst output of TRN neurons, however whether the contribution of SK-influence on burst discharge is heterogeneous throughout the TRN is unclear.

The demarcation between burst- and tonic-discharge of thalamic neurons has historically been dramatic based on the apparent all of none burst discharge of high frequency action potential discharge ^14^. In this study, we focused on the burst discharge of TRN neurons considering the downstream inhibitory impact on thalamic nuclei strongly impacts rhythmic activity within thalamocortical circuits. Surprising to us, burst discharge of TRN neurons spans a broad range of discharge frequencies. These experiments were designed to determine potential mechanisms underlying the heterogeneity of burst frequencies. Findings from this study are aimed to improve characterization of TRN function and classify how intrinsic physiology of these neurons relates to the diverse functions of the TRN.

## MATERIALS AND METHODS

### Slice preparation

All animals were housed at the Michigan State University animal facilities and experimental procedures were conducted in accordance with the Institutional Animal Care and Use Committee (IACUC). Adult female and male C57BL/6J (WT), and SOM-IRES-Cre (Jackson Laboratories #013044)^26^ x Ai14 (Jackson Laboratories #007914)^27^. mice were utilized within these experiments (postnatal age: 42 ± 12 days old, range 30 to 111 days old). Animals were deeply anesthetized (3% isoflurane, 97% O_2_) and in some cases perfused with cold (4°C) oxygenated (95% O_2_, 5% CO_2_) slicing solution containing (in mM): 2.5 KCl, 1.25 NaH_2_PO_4_, 10.0 MgSO_4_, 0.5 CaCl_2_, 26.0 NaHCO_3_, 10.0 glucose, and 234.0 sucrose prior to decapitation. Brains were quickly removed and submerged in cold (4°C), oxygenated (95% O_2_, 5% CO_2_) slicing solution. Horizontal brain slices (300 μm thickness) were made using a vibrating tissue slicer, hemisected and transferred to a warmed holding chamber (35°C) with oxygenated (95% O_2_, 5% CO_2_) artificial cerebrospinal fluid (ACSF) solution containing (in mM): 126.0 NaCl, 2.5 KCl, 1.25 NaH_2_PO_4_, 2.0 MgCl_2_, 2.0 CaCl_2_, 26.0 NaHCO_3_, and 10.0 glucose. After 30 minutes, the holding chamber temperature was returned to room temperature and slices incubated for a minimum of 60 minutes prior to recording.

### Electrophysiology

Borosilicate glass micropipettes (4-6 MΩ resistance) were used to perform whole-cell recordings. Micropipettes were filled with a solution of (in mM): 117 K-gluconate, 13 KCl, 1.0 MgCl_2_, 0.07 CaCl_2_, 0.1 EGTA, 10 HEPES, 2.0 Na2-ATP, and 0.4 Na-GTP. Experiments utilized a Zeiss microscope equipped with DIC optics (Thorndale, NY). TRN was located using a low power objective (5x), and individual neurons were identified using a high-power, water immersion objective (63x). Cells were recorded in a slice chamber with fresh oxygenated (95% O_2_, 5% CO_2_) ACSF (2.5–3 ml/min) maintained at 32°C. Following break-in, cells were allowed to stabilize for at least five minutes prior to starting experiments. Only cells with an input resistance greater than 100 MΩ, resting membrane potential more negative than −58 mV, lack of spontaneous action potential firing, and evoked action potential peak >0 mV were included for analyses. Data were obtained using a MultiClamp 700b amplifier, sampled at 20 kHz, and filtered at 10 kHz (Digidata 1440a, Molecular Devices, San Jose, CA). Reported voltages are corrected for an 8 mV liquid junction potential. PClamp software (Molecular Devices, San Jose, CA) was used for data acquisition and analysis.

Pharmacological agents were purchased from Tocris (Minneapolis, MN) or Sigma-Aldrich Co. (St. Louis, MO). Concentrated stock solutions were made with distilled water and brought to final concentration in ACSF prior to experimentation. All antagonists were bath applied for at least 10 minutes prior to commencing experiment. For experiments using agonists, these compounds were applied via syringe pump ^28^.

### Morphology

Soma volume was calculated from confocal z-stack images spanning the entire depth of the neuron. Alexa-594 (25μM) was included in the internal solution to visualize neurons, and neurons were allowed to fill for at least 30 minutes prior to image acquisition. Images were obtained using a 2-photon laser scanning microscope (Ultima, Bruker) with the Prairie View (Bruker) acquisition system. To calculate soma volume, the z-stacks were imported into FIJI (NIH open source) and voxels contained within the soma were counted by analyzing the Thresholded Voxels via the Voxel Counter plug-in (NIH open source).

### Statistical Analysis

Burst frequency was calculated from the average discharge frequency of action potentials in the initial burst response. The burst was produced by the lowest intensity current step that produced a short latency, transient action potential discharge of ≥2 action potentials. Action potential threshold was calculated as the voltage at which the rise of the action potential surpasses 10 mV/ms. Rheobase was measured as the lowest intensity current step (20 pA intervals, 1 s duration) to elicit an action potential from holding membrane potential of −60 mV. Half-width of the action potential was defined as the duration of the action potential at half amplitude, measured from the threshold. Input resistance was measured from the linear slope of a current-voltage response around rest (minimum −10 to +10pA, 10-20 pA increments, 1 s duration, measurements taken from the last 200ms). Population values are reported as mean ± standard deviation unless otherwise stated.

## RESULTS

Whole-cell recordings were obtained from 248 TRN neurons to assess burst firing characteristics. Neurons were held at −80 mV, and increasing depolarized current steps (20 pA increments, 1s duration) were delivered until a burst discharge was elicited. Burst frequency was calculated as the average frequency (1/inter-spike interval) of all action potentials (APs) contained in the burst. Considering many of the bursts recorded contained only two APs, we compared the frequency of the first inter-spike interval to the average frequency of all APs in the burst within individual neurons. We found a strong correlation of these two measures (r^2^=0.859, p<0.001, n=165, ANOVA). and utilized the average frequency of the entire burst for the remainder of the studies. As illustrated in figure 1, we observed a broad range of burst frequencies from 9 to 342 Hz (Fig. 1A, B). As illustrated in Fig. 1C, cells with burst frequencies of ≤100 Hz typically had fewer APs/burst (2.7 ± 1.0 APs/burst, n=100), and in many cases there were only two APs/burst (61/100 cells). While non-bursting neurons have been identified in TRN, these cells lack the voltage-dependent non-linearity of AP discharge as seen from the burst discharge of thalamic neurons ^9,14,29–32^. Two APs/burst were characterized by having a burst firing frequency at least twice as fast as discharge frequency in tonic mode. In contrast, in the population of neurons with burst averages >100 Hz, 88% (135/148 neurons), the burst consisted of three or more APs/ burst. Based on prior studies, we anticipated the overwhelming population of burst frequency would be skewed to higher frequencies (i.e., >200 Hz), but were surprised to see such a broad variability in burst frequency output of the TRN neurons ^8,9,13,31,32^.

**Figure 1:**
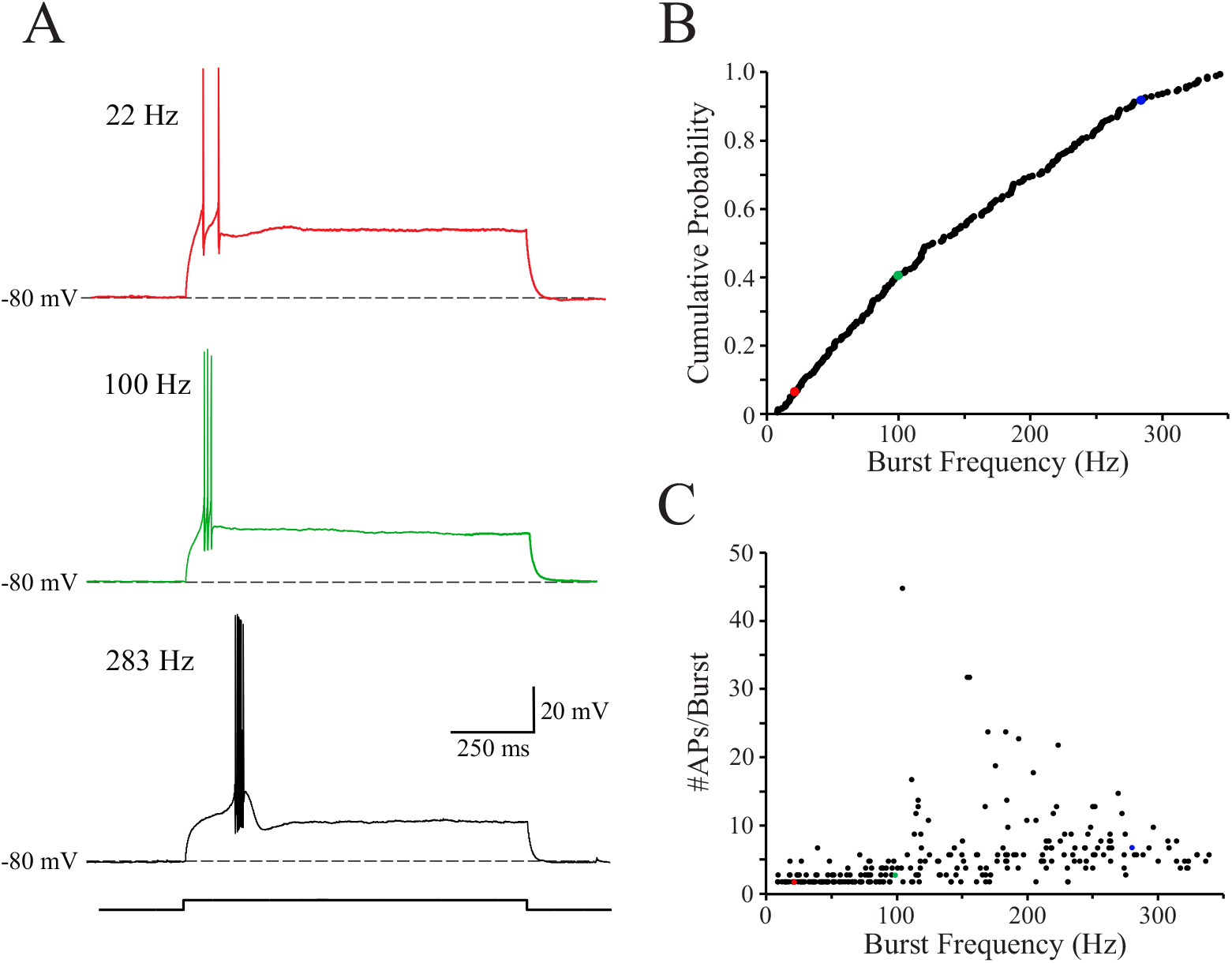
TRN neurons burst at a wide range of varying frequencies. **A.** Representative voltage traces of burst firing elicited from somatic current injection while held at Vm= −80 mV. Average burst frequency is shown in parentheses to the left or the trace. **B.** Cumulative probability burst frequency discharge of all TRN neurons recorded (n=248). **C.** Graph depicting the number of action potentials in the burst discharge as it relates to burst frequency.

### Synaptically evoked bursts do not have significantly increased frequencies compared to somatically evoked bursts

Prior studies indicate a wide distribution of T channels along the dendritic arbor of TRN neurons ^29,33^. If TRN neurons have more distally located T channels, perhaps the slower burst frequency could be due to incomplete activation of the dendritic T channels from the somatic current injection used to evoke the burst. Electrical stimulation of afferent synaptic inputs was utilized to provide activation of dendrites as opposed to somatic current injection. A stimulating electrode was placed within the internal capsule, and the stimulus intensity gradually increased (50-400 μA, 100 μs) to evoke a burst discharge. The burst frequency of the synaptically evoked burst was strongly correlated with the burst frequency of the somatically evoked burst produced by current step injection (Fig. 2A, B; r^2^=0.863, p<0.001, n=13). The number of APs/burst via either method was positively correlated and remained close to the unity line (Fig. 2 A, C; r^2^=0.567, p = 0.003, n=13), and there was no significant difference between the number of APs/burst (synaptic: 4.3 ± 3.0; somatic: 6.1 ± 5.8, n = 16, p=0.088, paired Student’s T test). Since no significant difference in burst firing frequency was observed between synaptic stimulation and somatic current injection this suggests the heterogeneity of the burst firing is intrinsic to the neurons and not a result of differential activation of soma versus dendrites.

**Figure 2:**
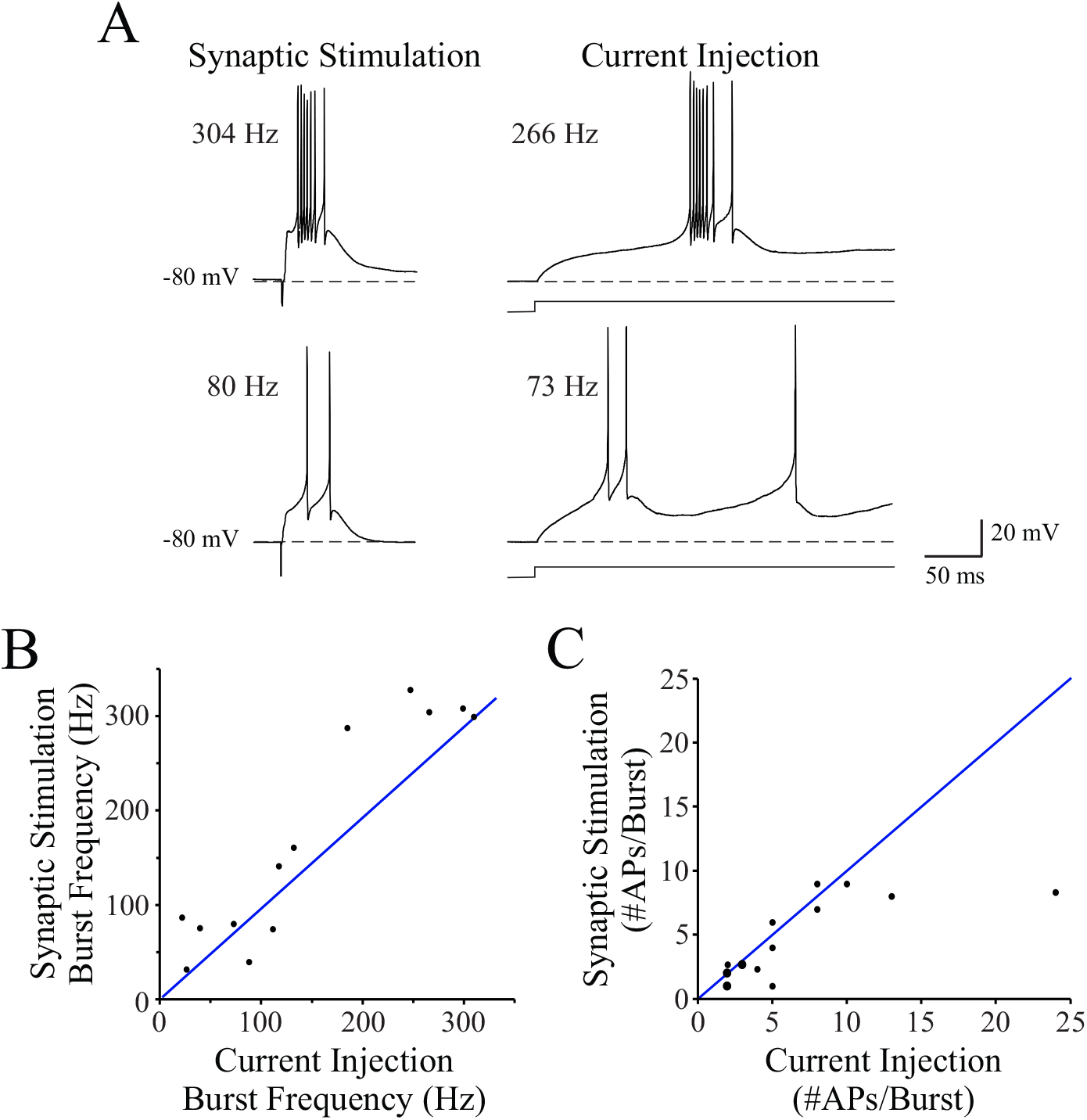
Comparison of synaptic vs. somatic activation of burst discharge in TRN neurons. **A.** Representative voltage traces of burst firing evoked by somatic current injection and electrical stimulation of synaptic afferents in a high frequency bursting (*top*) and low frequency bursting (*bottom*) neuron. Burst frequency for these individual traces is shown to the left of the response. **B.** Comparison of burst frequency produced by synaptic stimulation and somatic current injection (n=13). **C.** Number of APs/burst for synaptically evoked and somatically evoked burst discharges (n=16). Each of the three enlarged data points represent two identical overlapping data points.

### Intrinsic properties differ between higher and lower frequency bursting TRN neurons

We next determined if the heterogeneity of burst frequencies observed were related to differences in intrinsic properties of the TRN neurons. Input resistance and membrane potential were measured from rest, number of APs/burst and average burst frequency were measured from −80 mV, and AP half-width, threshold, and rheobase were measured from −60 mV to avoid activation of T channels. Initial correlation analyses revealed that several intrinsic properties were positively correlated with the average burst frequency (Table 1). The APs/burst (r^2^=0.168, p<0.001), tonic firing frequency (r^2^=0.136, p<0.001) and rheobase (r^2^=0.077, p=0.007), were positively correlated with average burst frequency. In contrast, the average burst frequency was negatively correlated with input resistance (r^2^=-0.036, p=0.004) and action potential half-width (r^2^=-0.150, p<0.001). There was no significant correlation of resting membrane potential (r^2^=0.009, p=0.143) or action potential threshold (r^2^=0.011, p=0.324) with the burst frequency.

**Table 1:**
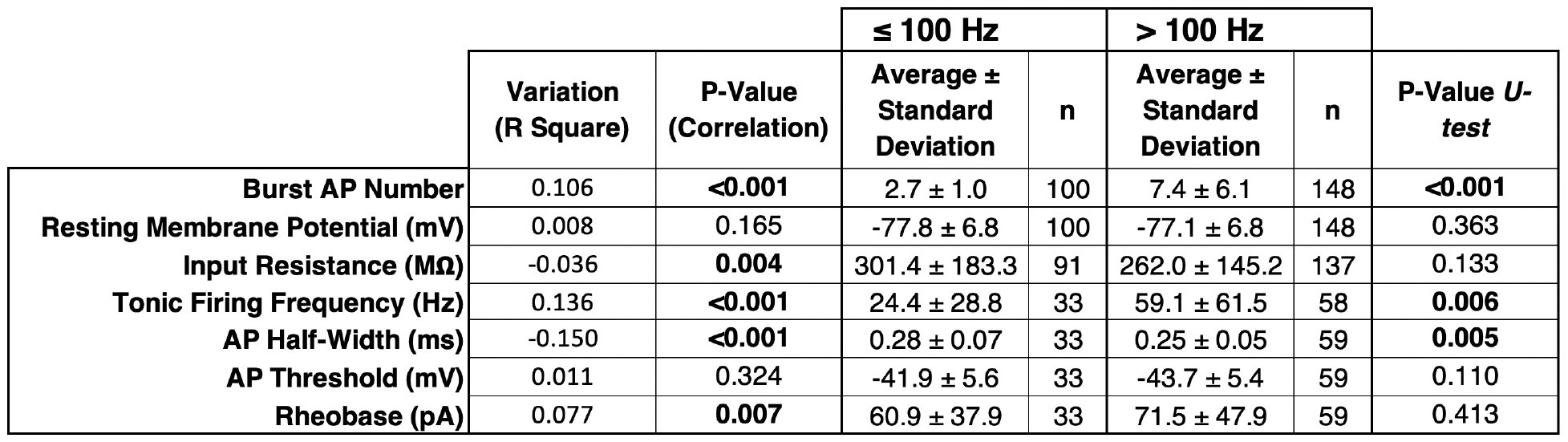
Intrinsic properties of TRN neurons.

|  |  |  | ≤ 100 Hz |  | > 100 Hz |  | P-Value *U*-test |
| --- | --- | --- | --- | --- | --- | --- | --- |
|  | Variation<br>(R Square) | P-Value<br>(Correlation) | Average ±<br>Standard<br>Deviation | n | Average ±<br>Standard<br>Deviation | n |  |
| **Burst AP Number** | 0.106 | **<0.001** | 2.7 ± 1.0 | 100 | 7.4 ± 6.1 | 148 | **<0.001** |
| **Resting Membrane Potential (mV)** | 0.008 | 0.165 | -77.8 ± 6.8 | 100 | -77.1 ± 6.8 | 148 | 0.363 |
| **Input Resistance (MΩ)** | -0.036 | **0.004** | 301.4 ± 183.3 | 91 | 262.0 ± 145.2 | 137 | 0.133 |
| **Tonic Firing Frequency (Hz)** | 0.136 | **<0.001** | 24.4 ± 28.8 | 33 | 59.1 ± 61.5 | 58 | **0.006** |
| **AP Half-Width (ms)** | -0.150 | **<0.001** | 0.28 ± 0.07 | 33 | 0.25 ± 0.05 | 59 | **0.005** |
| **AP Threshold (mV)** | 0.011 | 0.324 | -41.9 ± 5.6 | 33 | -43.7 ± 5.4 | 59 | 0.110 |
| **Rheobase (pA)** | 0.077 | **0.007** | 60.9 ± 37.9 | 33 | 71.5 ± 47.9 | 59 | 0.413 |

To further assess potential differences in intrinsic properties of low- and high-frequency burst TRN neurons, the neurons were divided into two groups: burst frequency ≤100 Hz or >100 Hz (Table 1). With this grouping, the number of APs/burst, tonic firing frequency, and AP half-width differed significantly. In contrast, resting membrane potential, input resistance, AP threshold, and rheobase did not significantly differ between the high and low frequency bursting neuron groups (Table 1).

### Lower frequency bursting neurons have smaller somata

Several studies suggest that TRN somatic morphology is not homogeneous and multiple subtypes can be differentiated based on cell morphology ^34–36^. In a subpopulation of TRN neurons (n=64), cells were filled with a fluorescent dye (Alexa 594, 25 μM) and soma volume calculated from maximum z-projections of the filled neurons. In this subset of neurons, the burst frequencies ranged from 9-325 Hz and the soma volumes were positively correlated with the burst frequency (Fig. 3A *left*; n=64, r^2^=0.22, p<0.001). The somas of the low frequency bursting neurons were significantly smaller than that of the high frequency bursting neurons (Fig. 3A *right*; ≤100 Hz: 1496 ± 615 μm^3^, n=23; >100 Hz: 1970 ± 796 μm^3^, n=41; p=0.016, Mann-Whitney U-test).

**Figure 3:**
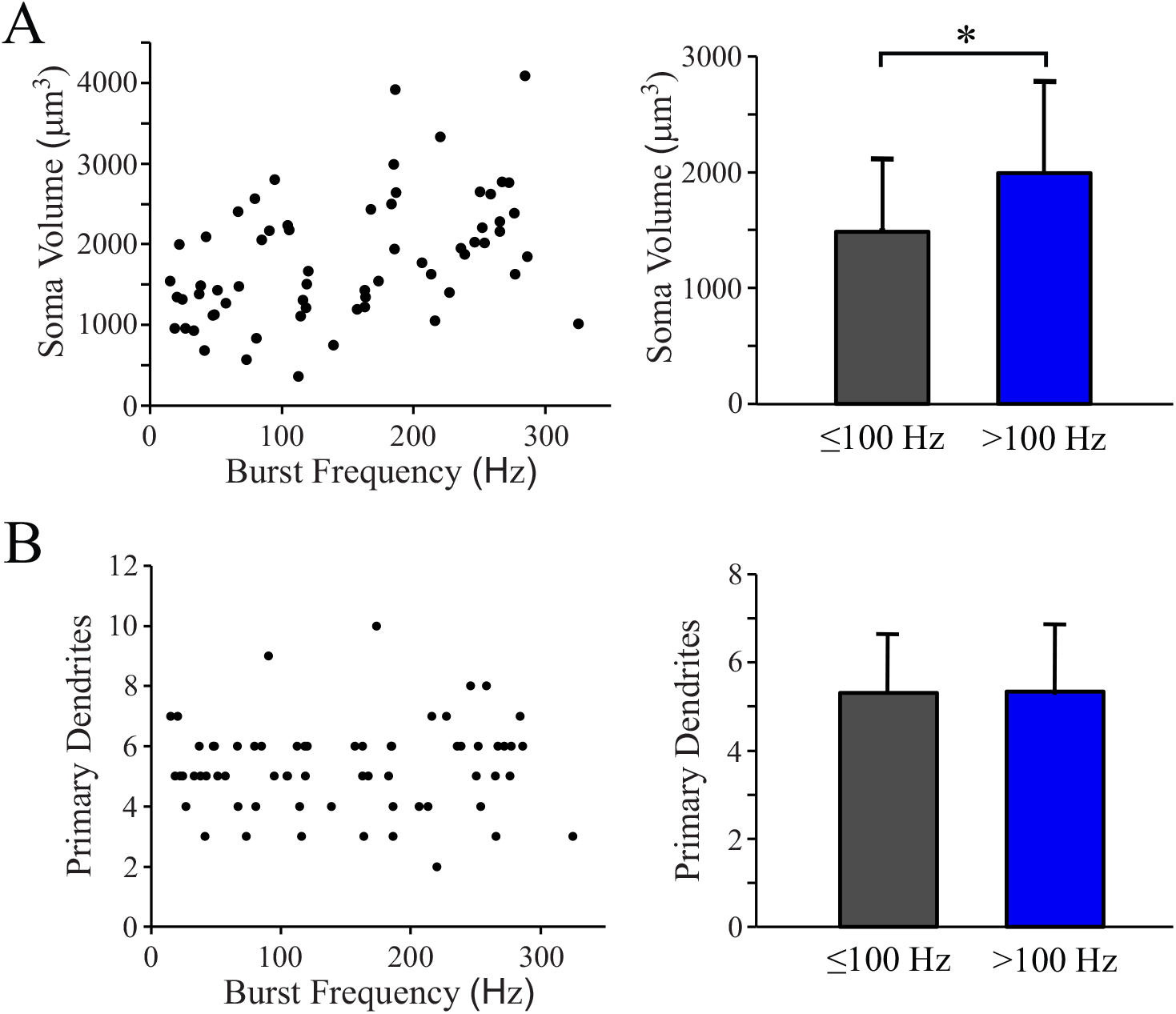
Morphologic comparison of low- and high-frequency bursting TRN neurons. **A.** *Left*, Population data of soma volume relative to burst frequency. *Right*, Histogram depicting soma volumes for ≤100 Hz and >100 Hz subgroups. **B.** *Left*, Population data depicting number of primary dendrites relative to to burst frequency. *Right*, Histogram depicting number of proximal dendritic branches for ≤100 Hz and >100 Hz subgroups. *, p < 0.05.

Previous studies have found that TRN neuron dendritic arborizations are heterogeneous with some cells having bipolar primary dendrites while others having multi-polar radial arborizations ^34,36^. Our findings indicate that there is no correlation between burst frequency and primary dendrite number (Fig. 3B *left*, n=64, r^2^=0.002, p=0.695) and no statistical difference in the number of primary dendrites between high frequency bursting neurons and low frequency bursting TRN neurons (Fig. 3B *right*, ≤100 Hz: 5.3 ± 1.3, n=23; >100 Hz: 5.3 ± 1.6, n=41, p=0.914, Mann-Whitney U-test).

### Burst frequency of TRN neurons is related to the underlying low threshold calcium response

In thalamic neurons, burst firing results from activation of T-type calcium channels giving rise to a transient depolarization (LTS: low threshold spike) upon which multiple APs can occur. To investigate the underlying depolarization producing the burst firing, tetrodotoxin (TTX) was added to the bath to block AP discharge. All neurons tested in 1 μM TTX (n=40) produced a transient depolarization in response to depolarizing current steps (Fig. 4A). The peak of the transient depolarization was positively correlated with the burst frequency discharge prior to TTX application (Fig. 4B *left*, r^2^=0.6504, p<0.001). The LTS amplitude was significantly larger in high burst frequency neurons compared to the low burst frequency neurons (Fig. 4B *right*; ≤100 Hz: 9.5 ± 6.2 mV, n=24; >100 Hz: 20.4 ± 8.8 mV, n=24; p<0.001, Mann-Whitney U-test). We next tested whether this smaller transient depolarization could be due to partial activation of the channels responsible for the depolarization. If the channels responsible were being only partially activated, the peak amplitude should increase with larger current injections. A series of increasing magnitude current steps (20 pA increments) were injected, and the peak of the transient depolarization amplitude was measured at each current step. Results revealed that for both higher and lower frequency bursting neurons, the transient depolarization did not increase in magnitude with increasing current injection for either low- or high-frequency bursting neurons (Fig. 4C: Low-frequency; F_4,60_=0.918, p=0.459. High-frequency; F_4,69_=2.451, p=0.0541, ANOVA).

**Figure 4:**
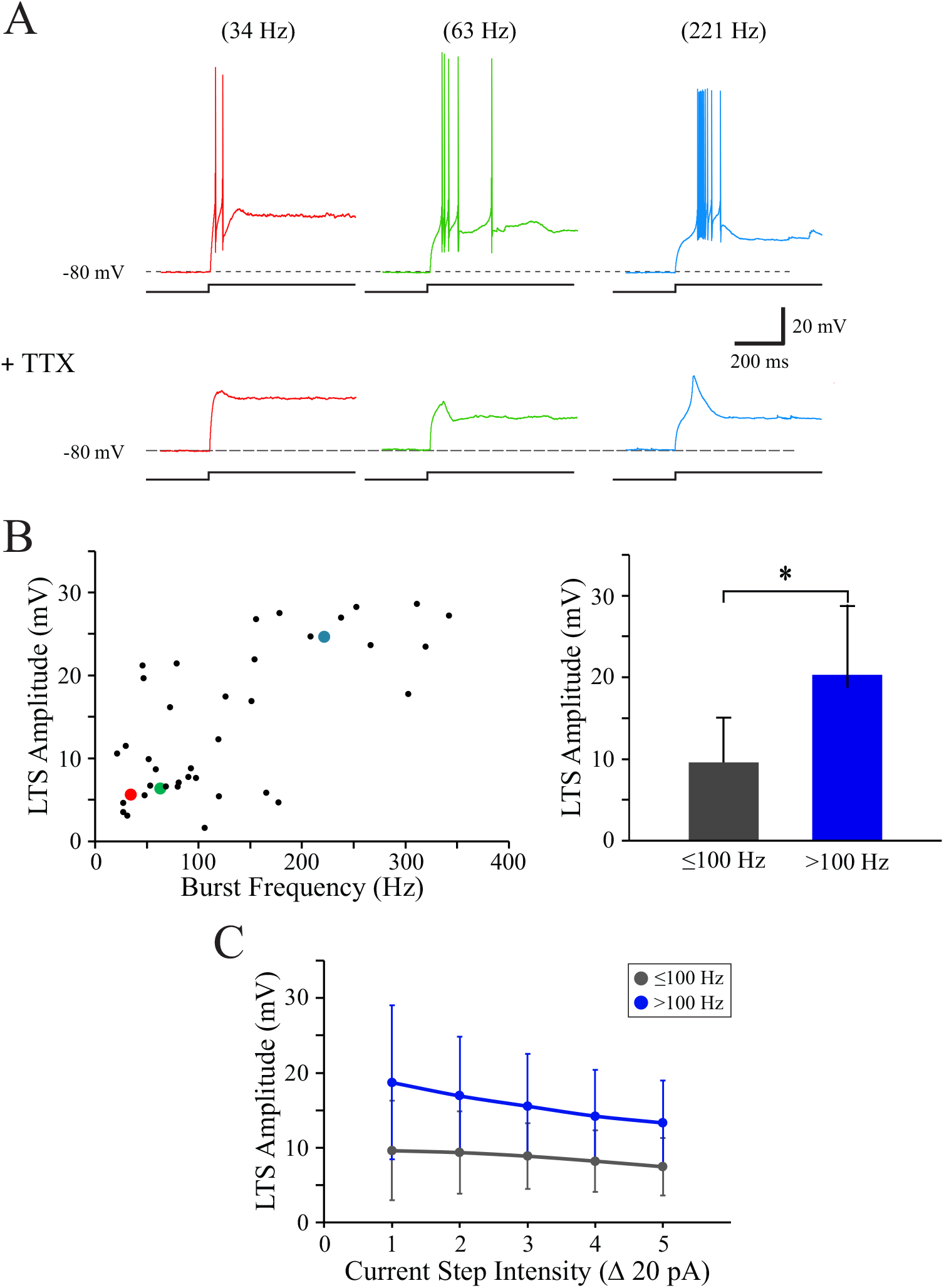
Transient depolarization amplitude is correlated with burst frequency. **A.** *Top*, Representative voltage traces of three TRN neurons with different burst frequencies (red: 34 Hz; green: 63 Hz; blue: 221 Hz). *Bottom*, Same current steps as **A** except in the presence of TTX (1 μM) to unmask the underlying transient depolarization (LTS). **B.** *Left*, Scatter plot of peak LTS amplitude (transient depolarization) relative to burst frequency of neuron prior to TTX exposure. Data points from the three example cells in **A** are designated by respective color-coded data points. *Right*, Summary histogram of LTS amplitude in low- (≤100 Hz, gray) and high-frequency (>100 Hz, blue) bursting neurons. **C.** LTS amplitude in response to increasing stimulus intensity (Δ 20 pA) for higher (blue) and lower frequency (gray) bursting neurons. *, p<0.05.

To measure the inward current underlying the transient depolarization in TRN neurons, an inactivation protocol was conducted in voltage clamp in the presence of 1 μM TTX (Fig. 5A). In all neurons tested (n=27), which ranged from burst frequencies of 17 to 326 Hz prior to the voltage clamp recordings in TTX, a transient inward current was evoked. The peak of the inward current was positively correlated with the burst frequency (Fig. 5B *left*, n=27, r^2^=0.780, p<0.001). The peak of the inward current in low frequency bursting neurons was significantly smaller compared to the current amplitude in the high frequency bursting neurons (Fig. 5B *right*; ≤100 Hz: 182.6 ± 77.6 pA, n=14; >100 Hz: 481.0 ± 196.7 pA, n=13; p<0.001, Mann-Whitney U-test) In addition, the half-width of the inward current was significantly longer in the low frequency bursting neurons (Fig. 5C; ≤100 Hz: 31.6 ± 13.6 ms, n=12; >100 Hz: 22.8 ± 11.0 ms, n=11; p=0.026, Mann-Whitney U-test). In a subgroup of neurons, both the transient depolarization (current clamp recording) and the transient inward current (voltage clamp recording) were recorded from the same neurons. In these cells, there was a positive correlation between the two transient responses (Fig. 5D, n=14, r^2^=0.812, p<0.001).

**Figure 5:**
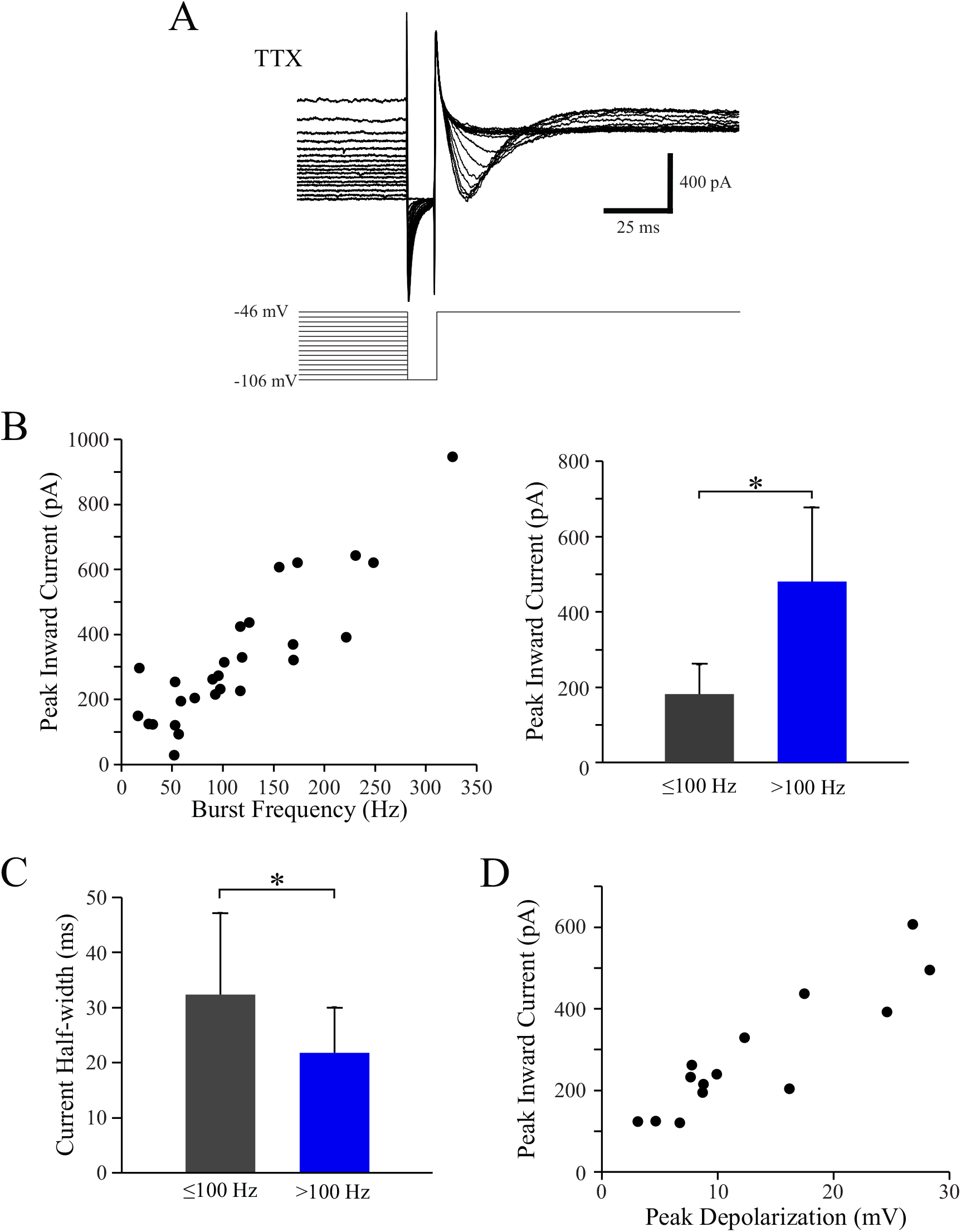
Transient inward current is correlated with burst frequency. **A.** Current traces from representative high frequency bursting neuron in response to inactivation voltage command protocol used to evoke transient inward current in present of TTX (1 μM). **B.** *Left*, Peak inward current relative to burst frequency of neuron prior to TTX exposure. *Right*, Summary data indicating peak inward current of low frequency (black) and high frequency (blue) bursting neurons. **C.** Summary data of inward current half-width in low- (black) and high-frequency (blue) bursting neurons. **D.** Peak inward current (voltage clamp recording) is positively correlated with the peak transient depolarization (current clamp recording). *, p<0.05.

### SK channels modulate burst characteristics in TRN neurons

Small-conductance calcium-activated potassium (SK) channels impact TRN neuron output by rapidly attenuating the burst discharge ^23,25,37,38^. We tested if blocking SK channels would differentially affect burst discharge in low- or high-frequency bursting TRN neurons. In the presence of apamin (100 nM), the APs/burst was significantly increased in both low frequency bursting neurons (Fig. 6A, D, control: 2.6 ± 1.2; apamin: 10.8 ± 5.9, n=11, p<0.001, paired Student’s T test) and high frequency (Fig. 6A, D, control: 6.5 ± 3.3, apamin: 25.8 ± 19.5, n=11, p=0.08, paired Student’s T test). The burst frequency was also significantly increased by apamin in low frequency bursting neurons (Fig. 6A, D, control: 42.7 ± 24.8 Hz, apamin: 177.6 ± 113.3 Hz, n=11, p=0.001, paired Student’s T test) and high frequency bursting neurons (Fig. 6A, D, control: 207.8 ± 59.9 Hz, apamin: 303.4 ± 104.3 Hz, n=11, p=0.002, paired Student’s T test).

**Figure 6:**
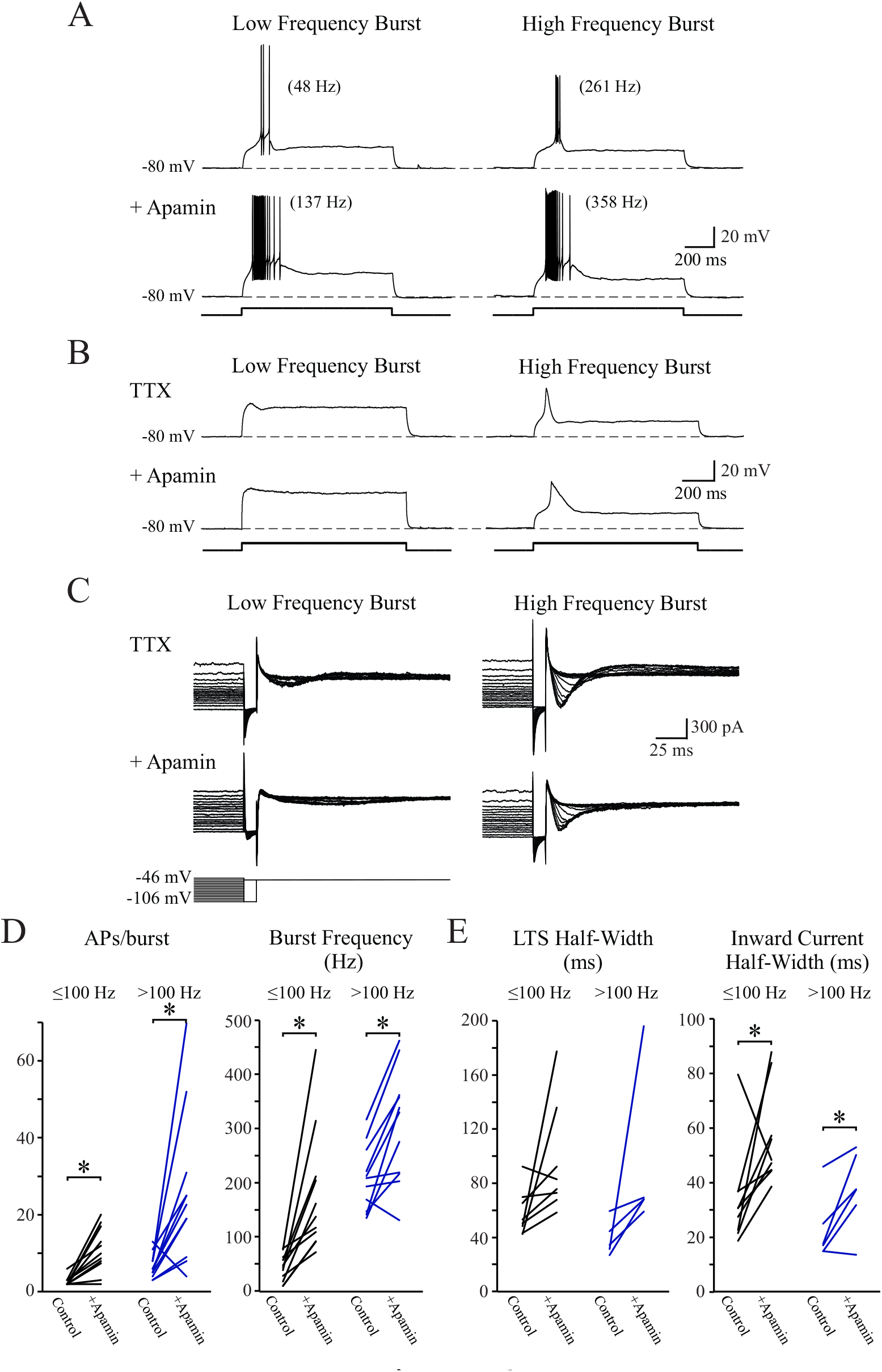
Apamin prolongs the individual burst discharge in TRN neurons: **A.** Representative voltage traces of a low frequency (48 Hz) and high frequency (261 Hz) bursting neuron before and after application of apamin (100 nM). **B**. Voltage recordings in the presence of TTX (top, 1 μM) revealing the underlying LTS from representative lower frequency bursting (78 Hz, pre TTX) and higher frequency bursting (252 Hz, pre TTX) neurons. Voltage traces from the same neurons as above in the presence of apamin (bottom, 100 nM). **C.** The same neurons recorded in **B**, in voltage clamp showing the transient inward current in the presence of TTX (top, 1 μM) and in the presence of apamin (bottom, 100 nM). **D.** In apamin, nearly all TRN neurons discharge more APs/burst compared to control. In addition, burst frequency also increases in apamin. Post apamin burst frequencies are calculated by same number of initial APs in the pre-apamin condition. **E**. Population data depicting the LTS half-width (*left*, current clamp recording) and inward current half-width (*right*, voltage clamp recording) for low- and high-frequency bursting neurons before and after application of apamin. *, p<0.05.

Considering the facilitation of burst activity following the attenuation of SK, we next tested if the underlying LTS was altered by blocking SK. In the presence of TTX (1 μM), the addition of apamin did not significantly alter LTS amplitude in low- or high-frequency burst neurons (Fig. 6B, ≤ 100 Hz: control: 8.2 ± 3.9 mV, apamin: 9.6 ± 4.2 mV, n=8, p=0.06, paired Student’s T test; > 100 Hz: control: 16.9 ± 9.7 mV, apamin: 18.9 ± 5.6 mV, n=6, p=0.19, paired Student’s T test). However, the LTS half-width, was increased in 7/8 lower frequency bursting neurons (Fig. 6B, E, control: 58.2 ± 16.9 ms, apamin: 95.4 ± 40.6 ms, n=8, paired Student’s T test, p=0.06), and all higher frequency bursting neurons (Fig. 6B, E, control: 39.3 ± 13.0 ms, apamin: 92.2 ± 58.3 ms, n=5, paired Student’s T test, p=0.136), these increases were statistically insignificant. It must be noted that the data is skewed because in two cells there was a long-lasting plateau potential that prevented measurement of the width.

We next examined the effect of apamin on the transient inward current. The peak amplitude of the inward current was not altered in the presence of apamin for either low frequency bursting neurons (Fig. 6C; control: 192.1 ± 60.5 pA, apamin: 188.0 ± 69.1 pA, n=9, p=0.77, paired Student’s T test) nor the high frequency bursting neurons (Fig. 6C, control: 464.2 ± 114.9 pA, apamin: 392.7 ± 118.0 pA, n=7, p=0.12, paired Student’s T test). The half-width of the transient inward currents were significantly increased by apamin in low frequency bursting neurons (Fig. 6C, E, control: 33.9 ± 18.3 ms, apamin: 56.5 ± 17.6 ms, n=9, p=0.03, paired Student’s T test) and high frequency bursting neurons (Fig. 6C, E, control: 20.8 ± 11.9 ms, apamin: 35.2 ± 14.0 ms, n=7, p=0.03, paired Student’s T test).

### Low-threshold depolarizations in both lower and higher frequency bursting neurons are calcium-mediated

We have found a transient low-threshold depolarization and associated inward current present in both low- and high-frequency bursting neurons. The inward current is significantly lengthened by blocking SK channels. To confirm that this current is calcium-mediated, we tested the sensitivity to 100 μM NiCl_2_ application, which has been shown to preferentially block T-type currents at relatively low concentrations ^13^. The transient inward current was measured from the maximum inward current in voltage clamp. The peak inward current was significantly attenuated by NiCl_2_ in all neurons tested (Fig. 7A, B; ≤100 Hz: pre-NiCl_2_: 190.3 ± 62.3 pA; NiCl_2_: 45.8 ± 37.5 pA, n=8, p<0.001, paired Student’s T test; >100 Hz: pre-NiCl_2_: 445.8 ± 217.5 pA; NiCl_2_: 178.0 ± 136.1 pA, n=9, p=0.003, paired Student’s T test). We also evaluated the effect of NiCl_2_ on the transient depolarization in current clamp recordings in the presence of TTX (1 μM). In low frequency bursting neurons, NiCl_2_ significantly attenuated the amplitude of the transient depolarization (Fig. 7C; pre-NiCl_2_: 6.4 ± 3.1 mV, NiCl_2_: 0.5 ± 0.9 mV, n=7, p=0.005, paired Student’s T test). Similarly, in high frequency bursting neurons, NiCl_2_ (100 μM) significantly attenuated the amplitude of the transient depolarization (Fig. 7C; pre-NiCl_2_: 22.1 ± 6.9 mV, NiCl_2_: 5.9 ± 6.6 mV, n=8, p<0.001, paired Student’s T test). These data indicate the calcium dependence of the transient depolarization in all TRN neurons tested.

**Figure 7:**
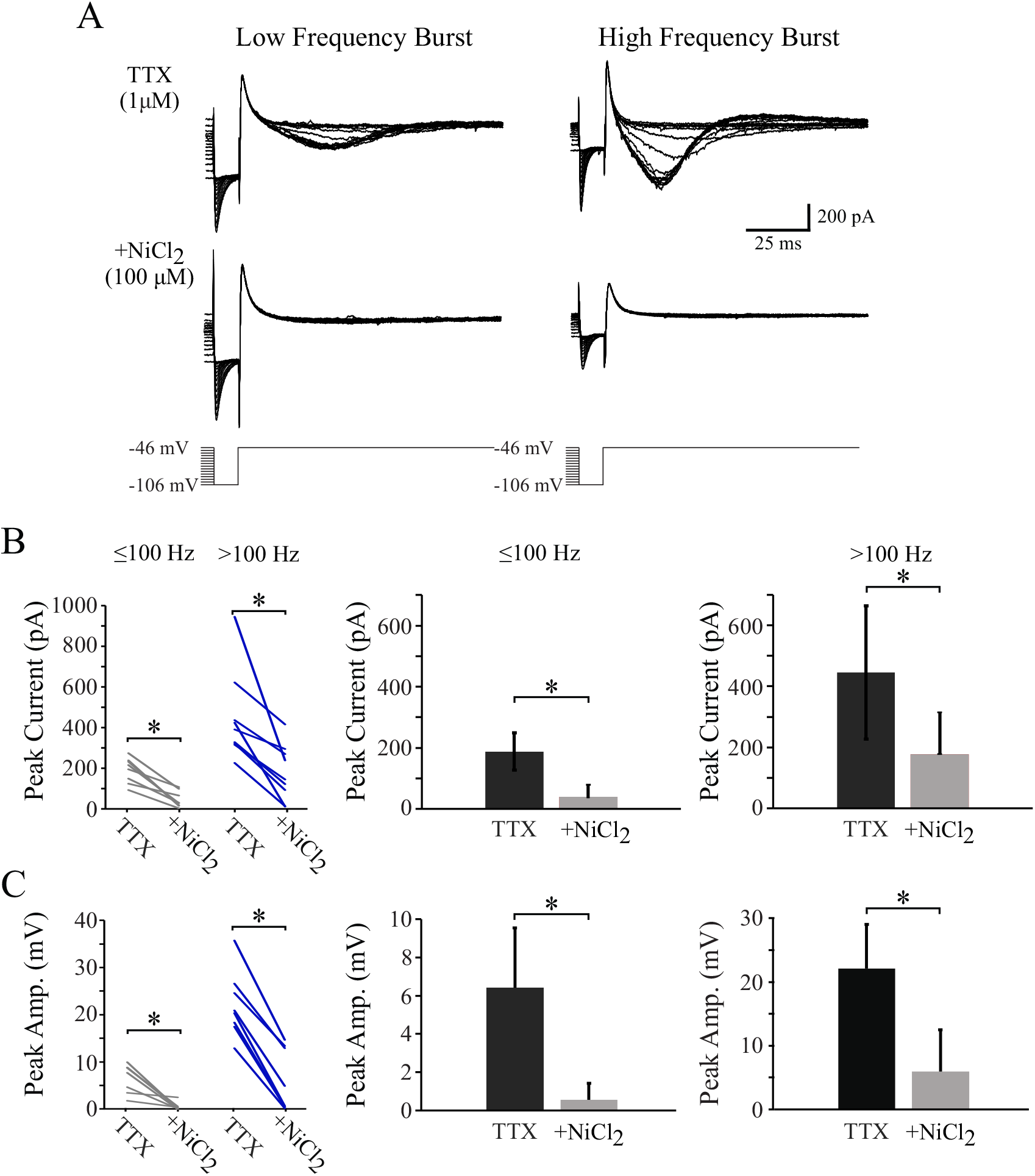
Nickel decreases the magnitude of transient inward current in TRN neurons. **A.** Representative transient inward current responses for a lower frequency (17 Hz) and high frequency (117 Hz) bursting neurons in TTX (1 μM), prior to and following the addition of NiCl_2_ (100 μM). **B.** *Left*, Population data indicating the impact of NiCl_2_ (100 μM) on the peak inward current in both low- (gray) and high-frequency (blue) bursting neurons. *Middle and right*, Bar graphs summarizing the attenuation of the inward current in both subpopulations of TRN neurons. C. Similar layout as **B**. Summary of the impact of NiCl_2_ (100 μM) on the transient depolarization (LTS) which was recorded in current clamp conditions. Similar to the transient inward current, the transient depolarization is significantly attenuated by NiCl_2_. *, p<0.05.

## DISCUSSION

In contrast to the traditional dogma that burst discharge is a high frequency transient discharge of many action potentials, we found that burst discharge of TRN neurons is quite distributed. Our data show that in mature mice, variable burst discharge frequency is related to multiple differences of intrinsic properties that may bring useful context to future studies of possible structural or functional subtypes. Neurons at either end of this variable burst frequency spectrum vary significantly in number of APs/burst, input resistance, and AP threshold and AP half-width. In addition to difference in intrinsic properties, low frequency bursting neurons have smaller LTS amplitudes, suggesting differences in T-channel number, location, or activation. In addition to T-channels, SK channels influence the frequency and duration, which likely influences downstream effects. Distinctions in these properties between neurons on either end of the burst frequency spectrum varied gradually, and there was no clear subtype grouping among TRN neurons based upon these described properties.

### Molecular markers of low-frequency bursting neurons

Recent genetic analysis performed on TRN neurons found that two identified gene profiles were expressed in a somewhat mutually exclusive manner ^39^. These genetic profiles included genes for GABA transporters (*Slc6a1*) and T-type calcium channels (*Cacna1i*), and as one profile increased in expression, the other appeared to decrease reciprocally. In consideration of our reported findings, it is possible that TRN neurons exhibit a spectrum of these described gene expression profiles, where neurons that highly express one profile have distinct electrophysiological properties from neurons with lower expression. Another recent study identified that TRN neurons located within the “core” and “edge” regions of the nucleus may be functionally distinct. Neurons at the edges of the nucleus tended to fire more lower frequency bursts than core cells. Calbindin-containing core cells received inputs from a first-order primary sensory relay, while the somatostatin (SOM)-containing edge cells received inputs from the higher-order nucleus ^40^. Given the previously reported connection between SOM-containing neurons and slower bursts ^30,40^, we recorded from SOM^+^ neurons within the SOM-Cre x Ai14 transgenic animal to determine whether lower frequency bursting neurons were SOM^+^. We recorded from 11 fluorescently-labeled SOM+ neurons from 3 mice and found an average burst frequency of 104 ± 99 Hz (4-339Hz), which overlapped with our non-labeled population (data not shown).

### Cell morphology and burst frequency

While soma size and volume has been previously measured in the TRN ^34^, to our knowledge it has never been associated with burst frequency. Our study found that mouse TRN somata volume ranged from 360-4090 μm^3^, whereas a previous study comparing soma volumes of rat, cat, and rabbit, reported were much smaller (150-860 μm^3^) ^34^. The findings that soma size varies with burst frequency supports the possibility of genetic differences among cells with different burst frequencies. Further studies investigating the afferent and efferent connections of slower and faster bursting neurons will clarify this question.

### SK channel contribution of burst discharge morphology

SK channel activation plays a critical role in shaping burst mode output in both low- and higher-frequency bursting neurons, and loss of I_SK_ interrupts TRN burst rhythms ^23,41^. I_SK_ is an important factor in the pattern of electrical output from a TRN neuron. We hypothesized that activation of I_SK_ may more strongly counter the effect of I_T_ in low frequency bursting neurons, resulting in a shorter and potentially weaker burst discharge. Greater I_SK_ will allow for multiple sequential bursts with shorter intervals in between. The converse would predictably result in longer bursts, potentially with more action potentials, and a longer refractory period. We consider that neurons with greater activity of I_SK_ may more likely participate in oscillatory burst rhythms, such as those seen in slow-wave sleep or epileptic seizures.

Another aspect to consider is that the location of the SK channels along the dendrite may have implications on the degree to which excitation from dendritic inputs (synaptic integration?) lead to downstream communication. Little is known about whether SK channels are distributed equally along the dendritic arbors of TRN neurons. A neuron with more distally located I_SK_ has the potential to discharge shortened the bursts form distal inputs, which are thought to be from an increased proportion of corticothalamic neurons ^42–44^. More proximal inputs on the same neuron, thought to have a higher representation from thalamocortical neurons, and would promote unaltered bursts that are longer and more robust. This scenario could allow TRN neurons to respond differently to the same magnitude of input depending on the originating location. Further studies into the location of SK channels on the dendrites will be important in determining location-specific impacts.

### Functional significance of heterogenous burst discharge

These studies revealed that neurons of the TRN cannot easily be separated into discrete populations based on electrophysiological or morphological properties. However, neurons at either end of this broad spectrum vary significantly in their intrinsic properties and the magnitude and duration of the burst-mode output. TRN neurons release GABA onto downstream thalamocortical neurons, which activate GABA_A_ and GABA_B_ receptors. In order to engage downstream metabotropic GABA_B_ receptors that elicit a longer lasting inhibitory drive, neurons must fire fast trains (> 100 Hz) of many action potentials ^20^. Lower frequency bursting neurons may be physiologically designed to only engage thalamocortical GABA_A_ receptors. In this way, higher and lower frequency bursting neurons may more finely modulate thalamic inhibition.

While high frequency bursting neurons appear necessary to maintain strong intrathalamic oscillations that are associated with sleep and certain pathophysiological conditions, the functional consequence of the low frequency bursting neurons is unclear. It is unlikely that these neurons play a significant role in the oscillatory activity in the thalamus given their lack of robust burst activity. One possible role of lower frequency bursting neurons could be to provide longer lasting, lower levels of thalamic inhibition to more finely modulate the circuits involved in sensory processing. While higher frequency bursting neurons are firing in a phasic “on or off” pattern, the lower frequency bursting neurons would act as a dimmer switch to maintain a lower level of inhibition that may more closely resemble a tonic or phasic pattern.

TRN neurons with burst frequencies at either end of the spectrum may have profoundly different impacts on thalamocortical inhibition of their downstream targets. These differences in frequency appear to be due to differing activity of dendritic T-type calcium channels, possibly due to altered levels of gene expression. Such differences in inhibitory output lead to both linear and non-linear voltage-dependent inhibition arising from the TRN. While neurons at either end of the spectrum have several intrinsic differences, these neurons do not appear to exist as separate groups, but as two ends of a complexly integrated spectrum of burst frequencies. In the TRN, bursting neurons are often studied as a homogenous cell type, however these studies have shown that bursts elicited from TRN neurons are highly variable and may act as uniquely designed processors for their specific synaptic inputs and thalamocortical target cells.

## Notes

Grants: This research was supported by NIH grant EY014024 and internal funds from Michigan State University.

### Competing Interest Statement

The authors have declared no competing interest.

